# The homoeologous genes for the Rec8-like meiotic cohesin in wheat: structure, function, and evolutionary implication

**DOI:** 10.1101/274522

**Authors:** Guojia Ma, Wei Zhang, Liwang Liu, Wun S. Chao, Yong Qiang Gu, Lili Qi, Steven S. Xu, Xiwen Cai

**Affiliations:** Department of Plant Sciences, North Dakota State University, Fargo, ND 58108, USA.; USDA-ARS, Red River Valley Agricultural Research Center, Fargo, ND 58102, USA.; USDA-ARS, Western Regional Research Center, Albany, CA 94710, USA.

**Keywords:** Rec8 cohesin, gene cloning, homoeoallele, meiosis, polyploid, and wheat

## Abstract

The Rec8-like cohesin is a cohesion protein essential for orderly chromosome segregation in meiosis. Here, we cloned two *Rec8*-like homoeologous genes (homoeoalleles) from tetraploid wheat (*TtRec8-A1* and *TtRec8-B1*) and one from hexaploid wheat (*TaRec8-D1*), and performed expression and functional analyses of the homoeoalleles. Also, we identified other two *Rec8* homoeoalleles in hexaploid wheat (*TaRec8-A1* and *TaRec8-B1*) and the one in *Aegilops tauschii* (*AetRec8-D1*) by comparative analysis. The coding DNA sequences (CDS) of these six *Rec8* homoeoalleles are all 1,827 bp in length, encoding 608 amino acids. They differed from each other primarily in introns although single nucleotide polymorphisms were detected in CDS. Substantial difference was observed between the homoeoalleles from the subgenome B (*TtRec8-B1* and *TaRec8-B1*) and those from the subgenomes A and D (*TtRec8-A1, TaRec8-A1*, and *TaRec8-D1*). *TtRec8-A1* expressed dominantly over *TtRec8-B1*, but comparably to *TaRec8-D1*. Therefore, the *Rec8* homoeoalleles from the subgenomes A and D may be functionally more active than the one from the subgenome B in wheat. The structural variation and differential expression of the *Rec8* homoeoalleles indicate a unique cross-genome coordination of the homoeologous genes in the polyploid, and imply the distinction of the wheat subgenome B from other subgenomes in the origin and evolution.

**HIGHLIGHT:** This work revealed the structural and expression patterns of the *Rec8*-like homoeologous genes in polyploid wheat, implying a unique origin and evolutionary route of the wheat B subgenome.

## INTRODUCTION

Meiosis is a specialized cell division with one round of DNA/chromosome replication and two successive divisions of the nucleus, producing haploid gametes (i.e. egg and sperms) for sexual reproduction. The first meiotic division (meiosis I) allows maternal and paternal homologous chromosomes to pair, recombine, and segregate, and consequently reduces chromosome number by half. The second meiotic division (meiosis II), similar to mitosis, allows sister chromatids (i.e. two longitudinal subunits of a replicated chromosome) to segregate. The outcome of meiosis is four haploid daughter cells that eventually develop into gametes (Kleckner, 1996; Roeder, 1997). Fertilization of haploid male and female gametes restores chromosomes to the parental ploidy level in the offspring. Therefore, meiosis ensures genetic integrity and generates genetic variability as well over the sexual reproduction cycle. It governs the transmission of genetic materials and provides the cytological basis of heredity.

Meiotic chromosome segregation is coordinated by the orientation of sister kinetochores where spindle microtubules attach on a chromosome, and the cohesion protein complex called cohesins. Cohesins glue sister chromatids together prior to chromosome segregation at meiosis I and II (Watanabe, 2012). A variety of meiotic genes/proteins have been identified as essential for proper chromosome cohesion and kinetochore orientation in meiosis (Ishiguro and Watanabe, 2007). Among those, Rec8 cohesin, which is highly conserved in eukaryotes, plays a central role in chromosome cohesion and sister kinetochore orientation at meiosis I in model species (Watanabe and Nurse, 1999; Yu and Dawe, 2000; Chelysheva *et al.*, 2005). Several other meiotic proteins have been found to interact with Rec8, and coordinate the function of Rec8 as a meiotic cohesin and a regulator for kinetochore orientation (Watanabe, 2004, 2005; Yokobayashi and Watanabe, 2005). The Rec8 cohesin appears at the pre-meiotic S phase, and associates with sister chromatids in the centromeric region as well as along chromatid arms. Prior to anaphase of meiosis I, Rec8 on chromatid arms is cleaved by separase to facilitate resolution of chiasmata and segregation of paired homologues. However, Rec8 in the centromeric region persists till anaphase II. The centromeric Rec8 is cleaved by separase to ensure segregation of sister chromatids prior to anaphase II (Lee *et al.*, 2005; Watanabe, 2005). *Rec8*-like genes have been cloned and characterized in diploid plant models, including Arabidopsis (*Syn1*) (Bai *et al.*, 1999;Bhatt *et al.*, 1999; Cai *et al.*, 2003), maize (*afd1*) (Golubovskaya *et al.*, 2006), and rice (*OsRad21-4*) (Zhang *et al.*, 2006). However, knowledge of the *Rec8*-like gene and cohesin is very limited in polyploids.

Wheat, including hexaploid bread wheat (*Triticum aestivum* L., 2n=6x=42, genome AABBDD) and tetraploid pasta wheat (*T. turgidum* L. ssp. *durum*, 2n=4x=28, genome AABB), contains genetically related subgenomes (i.e. A, B, and D in bread wheat, and A and B in pasta wheat). It has large chromosomes and a variety of cytogenetic stocks, providing advantages for direct visualization and characterization of chromosomes and subcellular structures. However, its large (∼16 Gb) and complex allopolyploid genome make gene cloning and functional analyses a challenging task especially for the genes involved in the subcellular processes, such as meiosis and mitosis. In this study, we aimed to clone the subgenome-specific homoeologous genes (homoeoalleles) for the Rec8-like cohesin in polyploid wheat, and to characterize their gene structure and function at the molecular and subcellular levels.

## MATERIALS AND METHODS

### Plant materials, crosses, DNA extraction, and male meiocyte collection

Tetraploid wheat ‘Langdon’ (LDN) (*T. turgidum* ssp. *durum*, 2n=4x=28, genome AABB), hexaploid wheat ‘Chinese Spring’ (CS) (*T. aestivum*, 2n=6x=42, genome AABBDD), CS nulli-tetrasomic lines, and LDN D-genome disomic substitution lines (LDN DS) (Joppa and Williams, 1988) were used in the cloning and functional analysis of wheat *Rec8*-like genes. The diploid ancestors of the wheat A genome [*T. urartu* (PI 428213)] and D genome [*Aegilops tauschii* (RL5286)] were included in the expression analysis.

LDN, LDN 1D(1A), and LDN 1D(1B) were used as females to cross with *Ae. tauschii* RL5286. Hybrid plants were obtained from the crosses by immature embryo culture on MS medium (Murashige and Skoog, 1962) as described by Cai *et al.* (2010). LDN haploid plants were produced by pollinating LDN plants with fresh pollen from corn (*Zea mays* L.), and subsequent embryo culture following the procedure of Cai *et al.* (2010).

All plant materials were grown in the temperature-controlled greenhouse for sampling of male meiocytes, leaf, and root tissues. Total genomic DNA of the plant materials was extracted from leaf tissues as described by Faris *et al.* (2000). Meiotically staged male meiocytes (anthers) were sampled following the procedure of Cai (1994). Anther samples were stored either in liquid nitrogen for real-time PCR and Western blotting or in 1×Buffer A (Bass *et al.*, 1997) for the *in situ* immunolocalization experiments.

### RNA extraction and cDNA preparation

Total RNA was extracted from leaves, roots, and meiotically staged anthers using RNAqueous^®^-4PCR Kit (Life Technologies, Grand Island, NY, USA). Prior to cDNA synthesis, total RNA was treated with DNase I and purified with RNAqueous^®^-4PCR Kit (Life Technologies, Grand Island, NY, USA). After quantification with NanoDrop ND-100 spectrophotometer (Thermo Fisher Scientific Inc., Wilmington, DE, USA) and agarose gel electrophoresis, 1 µg of total RNA was used as template for cDNA synthesis with SuperScript III First-Strand Synthesis System (Invitrogen Corporation, Carlsbad, CA, USA).

### cDNA cloning of the Rec8-like gene in LDN

Meiotic cohesin Rec8 is highly conserved across eukaryotes, particularly in the grass family. The amino acid sequence of rice Rad21/Rec8-like protein Os05g0580500 (GenBank accession NP_001056426.1) was used as query for tBLASTn search against the wheat expressed sequence tags (ESTs) database (https://wheat.pw.usda.gov/wEST/blast/). The ESTs having high sequence homology with rice Rad21/Rec8-like protein were annotated based on the information of the *Rec8*-like genes available in model species and plants. Gene-specific primers were designed from the nucleotide sequences of the candidate ESTs using Primer3 (http://frodo.wi.mit.edu/), and were used to synthesize cDNAs from total RNA of the LDN anthers at early prophase I/pachytene stages. The 3’-and 5’-RACE (rapid amplification of cDNA ends) were performed to extend the cDNA sequence of the candidate gene in tetraploid wheat LDN. The final complete cDNA sequence of the candidate gene was amplified by the primer pair GM067F/GM065R (Table S2) that spans the start and stop codons of the gene, and cloned as described by Ma *et al.* (2006).

### Chromosomal localization of the Rec8-like gene in wheat

The gene-specific primer pair GM008F/GM008R was used to amplify the gene-specific genomic segments in the 21 CS nulli-tetrasomic lines and 14 LDN DS (Table S2). After chloroform purification and ethanol precipitation, the amplicons were visualized by cleaved amplified polymorphic sequence (CAPS) with the restriction enzyme *Dde*I on the denaturing polyacrylamide gel (Konieczny and Ausubel, 1993).

### Quantitative real-time PCR

Real-time RT PCR was performed to quantify relative transcript levels of the *Rec8-*like gene in leaves, roots, and meiotically staged anthers in LDN and its cytogenetic stocks using a 7300 Real-Time PCR System (Applied Biosystems, Foster City, CA, USA) as described by Chao (2008). The minimum number of highly synchronized anthers (4-10) at each of the meiotic stages were sampled for RNA extraction in the real-time RT PCR analysis. The *Rec8-*like gene specific primer pair GM010F/GM010R (Table S2) was used in the real-time RT PCR. The 18S rRNA gene-specific primer pair GM003/GM004 (Table S2) was selected to amplify the endogenous control for the quantitative PCR according to the dissociation test and primer validation. The 18S rRNA gene was verified to be stably expressed in our experimental conditions and was used as an endogenous reference gene in this study. Two technical and three biological replications were implemented in these experiments. The comparative C_*T*_ method was used to determine the transcript levels of the *Rec8-*like gene in different samples (test) relative to the anthers at interphase stage (control) as described by Chao and Serpe (2010). Fold difference in gene expression of test *vs.* control sample is 2^−ΔΔCT^, where ΔΔC_*T*_ = ΔC_*T,test*_ - ΔC_*T,control*_. ΔC_*T,test*_ is the C_*T*_ value of the test sample normalized to the endogenous reference gene, and ΔC_*T,control*_ is the C_*T*_ value of the control normalized to the same endogenous reference gene. The C_*T*_ value of each sample is the average of two technical replicates. C_*T*_ values from three biological replicates were averaged, and data of the anther samples at interphase were used as baseline expression (control).

### Differential expression analysis of the wheat Rec8 homoeoalleles

The transcripts of wheat *Rec8* homoeoalleles were differentially analyzed in LDN, LDN 1D(1A), and LDN 1D(1B), and their hybrids with *Ae. tauschii* by semi-thermal asymmetric reverse PCR (STARP) (Qi *et al.*, 2015; Long *et al.*, 2017). This special PCR technique converts single nucleotide polymorphisms (SNPs) into length polymorphisms by adding an oligonucleotide (5’-ACGAC-3’ or 5’-ATGAC-3’) to the 5’ end of the allele-specific primer. Three primers, including two-tailed allele specific forward primers (AS-primers F1 and F2) and one common reverse primer, were used to amplify the SNP alleles (Table S2). The 5’ ends of GM157F1 (AS-F1) and GM157F2 (AS-F2) have a M13 tail (5’-CACGACGTTGTAAAACGAC-3’). The M13-tailed primer with an attached fluorescence tag at the 5’ terminus was added to the GM157F1/F2/R PCR mixture for amplicon visualization in the IR2 4300 DNA Analyzer (Li-Cor, Lincoln, NE, USA). The 5’ ends of GM178F1 (AS-F1) and GM178F2 (AS-F2) had a different tail (5’-GCAACAGGAACCAGCTATGAC-3’) due to the fluorescence primer change (fluorescence-M13 primer to fluorescence-PEA primer) in the lab. In the PCR involving GM178F1/F2/R, a universal priming-element-adjustable primer (PEA-primer 5’-ATAGCTGG-Sp9-GCAACAGGAACCAGCTATGAC-3’) with an attached fluorescence tag at the 5’ terminus was added for amplicon visualization in the IR2 4300 DNA Analyzer (Li-Cor, Lincoln, NE, USA). The F1/F2/R primer combinations were tested for amplification efficiency in the corresponding genomic DNA. Further, the F1/R and F2/R primer pairs were tested separately in the critical DNA samples for allele differentiation (Qi *et al.*, 2015; Long *et al.*, 2017).

### Antibody production and affinity purification

A 464-bp cDNA segment of the *Rec8-*like gene in LDN was chosen, based on its hydrophobicity and sequence uniqueness, to raise antibody against the wheat Rec8-like cohesion protein. The cDNA segment was PCR amplified using the primer pair GM026F/GM026R. A restriction enzyme recognition site for *Eco*RI and *Sal*I was added to the 5’ ends of the forward and reverse primer, respectively, for cloning purpose. Three nucleotides (TCA) were added to the 3’ end of the *Sal*I recognition site within GM026R to generate a stop codon (TGA) in the expression constructs (Table S2). The amplicon was cloned into two expression plasmid vectors pGEX-4T-1 (Amersham Biosciences, Piscataway, NJ, USA) and pMAL-c2X (New England Biolabs, Ipswich, MA, USA), leading to two distinct constructs (*pGEX-R26* and *pMAL-R26*). After verified by sequencing, these two constructs were transformed into *E. coli* strain BL21-Star (DE3) (Invitrogen Corporation, Grand Island, NY, USA) for fusion protein induction as described by Chao *et al.* (2007). Upon IPTG (isopropyl β-D-1-thiogalactopyranoside) induction, the fusion polypeptides, pGEX-R26 and pMAL-R26, were accumulated as insoluble pellets and resolubilized after sonication. The total proteins were separated on the SDS-PAGE gel and the candidate bands were cut out as per the estimated molecular weight. Upon the verification with a protein identification test performed in the Vincent Coates Foundation Mass Spectrometry Laboratory at Stanford University (Stanford, CA, USA) (Figure S5), the polypeptide pGEX-R26 was used for immunization and generation of the polyclonal antibody in rabbits by Affinity BioReagents (ABR, Golden, CO, USA; now Thermo Fisher Scientific, Inc.).

The anti-pGEX-R26 crude serum was affinity-purified as described by Chao *et al.* (2007) with minor modifications. The affinity-purified pMAL-R26 polypeptide was first coupled to AminoLink coupling resin using the AminoLink Plus Immobilization Kit (Pierce Biotechnology, Rockford, IL, USA) and then incubated with the crude serum. After incubation, the mixture of crude serum and resin was loaded to the column for antibody isolation. The anti-Rec8 antibody was eluted with the IgG Elution Buffer (Pierce Biotechnology, Rockford, IL, USA) after washing the column with 20 column volumes of 1× Phosphate Buffered Saline (PBS; 137 mM NaCl, 10 mM PO ^3-^, and 2.7 mM KCl; pH 7.4). Aliquots of anti-Rec8 antibody were stored in the −80°C freezer for subsequent uses.

### Immunoprecipitation, Western blotting, and in situ immunolocalization

Protein extraction, immunoprecipitation, and Western blotting were performed following the procedures of Chao *et al.* (2007). About 400 mg anthers at each of the four distinct meiotic stages (interphase, early prophase I, metaphase I/anaphase I, and metaphase II/anaphase II) were collected for protein extraction in the Western blotting analysis. The anti-Rec8 antibody was diluted in a ratio of 1:500 for Western blotting. *In situ* immunolocalization was conducted to examine the endogenous Rec8 protein on wheat chromosomes at different meiotic stages as described by Golubovskaya *et al.* (2006) with minor modifications. The primary anti-Rec8 antibody was probed by the secondary Anti-Rabbit IgG (whole molecule)-FITC Antibody produced in goat (Sigma-Aldrich Co., St Louis, MO, USA), and chromosomes were counterstained by propidium iodide (PI). Two negative control experiments were performed to monitor the specificity of the antibodies in meiocytes. In the first negative control, the thin layer of polyacrylamide gel containing meiocytes was directly incubated with the secondary antibody. In the second one, the thin layer of polyacrylamide gel containing meiocytes was incubated with the primary anti-Rec8 antibody that was preabsorbed with the fusion polypeptide pGEX-R26.

### Microscopy

Microscopy was conducted using a Zeiss Axioplan 2 Imaging Research Microscope equipped with ApoTome component (Carl Zeiss Light Microscopy, Jena, Germany). Two-dimensional (2-D) and three-dimensional (3-D) images were captured and analyzed using Zeiss Axio Vision 4 software as described by Cai *et al.* (2010).

### Genomic DNA cloning and DNA/protein sequence analysis

Three cDNA fragments of the *Rec8*-like gene cloned in LDN were bulked as a probe to screen LDN BAC library as described by Huo *et al.* (2006) (Table S2). The positive BAC clones were further verified by PCR with the wheat *Rec8* gene-specific primers. The verified BAC clones were characterized by fingerprinting with *Hin*dIII and CAPS to identify the positive BAC clones that contain different homoeoalleles of the *Rec8* gene in LDN. Sub-cloning was performed to delineate the homoeoalleles into smaller genomic fragments for sequencing using the pWEB-TNC^(tm)^ Cosmid Cloning Kit (Epicentre Biotechnologies, Madison, WI, USA). The full-length genomic sequences of the *Rec8* homoeoalleles in LDN were obtained using the DNA Walking *SpeedUp*^TM^ Premix Kit II (Seegene, Inc., Gaithersburg, MD, USA).

The DNA and protein sequences of the *Rec8*-like gene in LDN were analyzed using BLASTP 2.2.26+ in the NCBI non-redundant database (www.ncbi.nlm.nih.gov). PEST motif was predicted using EPESTFIND (www.emboss.bioinformatics.nl/cgi-bin/emboss/epestfind). Other motifs were identified with Motif Scan (http://myhits.isb-sib.ch/cgi-bin/motif_scan). The gene structures of the *Rec8* homoeoalleles were analyzed and visualized using Splign (http://www.ncbi.nlm.nih.gov/sutils/splign/splign.cgi) and GSDS 2.0 software (Hu *et al.*, 2015). Comparative analysis was performed for the cDNA and predicted protein sequences of the *Rec8* homoeoalleles (*TtRec8-A1, TtRec8-B1*, and *TaRec8-D1*) using MultiAlin (http://multalin.toulouse.inra.fr/multalin/) and Clustal Omega (https://www.ebi.ac.uk/Tools/msa/clustalo/), respectively.

### Phylogenetic analysis

Comparative analysis of the Rec8-like cohesin orthologues from different species was performed using ClustalX 2.1. Bootstrap Neighbor-Joining phylogenetic tree was built with 1,000 bootstrap replications and Poisson model using MEGA 6.0, (Tamura *et al.*, 2013).

## RESULTS

### Cloning of the Rec8-like gene in tetraploid wheat

The Rec8-like cohesin is highly conserved in the DNA/protein sequences and subcellular function across eukaryotes (Watanabe and Nurse, 1999; T óth *et al.*, 2000; Chelysheva *et al.*, 2005; Golubovskaya *et al.*, 2006; Shao *et al.*, 2011). Rice (*Oryza sativa*) is a model species closely related to wheat (Kellogg, 2001; Gaut, 2002; Salse *et al.*, 2008). In this study, we used the amino acid sequence (608 AA) of rice Rad21/Rec8-like protein Os05g0580500 (previous GenBank accession NP_001056426.1; now BAF18340.1) as query to search against the wheat EST database [wEST peptide pool (aa)] by BLASTp at the GrainGenes website (https://wheat.pw.usda.gov/wEST/blast/). An 808-bp wheat EST (GenBank accession BQ744508) was found to have 80% identity with the query at an E-value cutoff of e^−110^. This EST was annotated as part of the candidate gene for the Rec8-like cohesin in wheat.

Gene-specific primers were designed based on the DNA sequences in the conserved regions of the candidate EST. RT-PCR was performed with the cDNAs of LDN anthers at early prophase I through pachytene stages in which the *Rec8*-like genes were highly expressed according to the findings in model species (Lee *et al.*, 2005; Watanabe, 2005). The amplicons of RT-PCR were sequenced and analyzed, and then used to design new gene-specific primers for rapid amplification of cDNA ends (RACE). The cDNA assemblies of the candidate gene were obtained after several rounds of RACE in LDN. Since LDN is an allotetraploid with two homoeologous subgenomes (A and B), it generally contains two homoeoalleles with high sequence similarities on each of two homoeologous chromosomes (Murai *et al.*, 1999; Huang *et al.*, 2002; Kimbara *et al.*, 2004; Zhang *et al.*, 2011; Brenchley *et al.*, 2012). To eliminate assembling errors of the RACE segments, the gene-specific primers for the final round of RACE at 5’ and 3’ ends, which spanned the start and stop codons, were used to amplify the full-length coding DNA sequence (CDS) of the candidate homoeologous genes for the Rec8-like cohesin in LDN. Only one CDS, instead of two, was recovered and cloned from LDN. It was 1,827 bp in length encoding 608 amino acids with a predicted molecular weight of 67.6 kDa.

The alignments of the predicted protein for the candidate gene with the Rec8 orthologues from other eukaryotes revealed high levels of amino acid sequence similarity, especially with the monocots *Brachypodium distachyon*, rice, and maize (67-80%) (Table S1). In addition, the predicted protein of the candidate gene contains two conserved domains of Rad21/Rec8 cohesin, pfam04825 at N-terminus and pfam04824 at C-terminus. Also, the predicted protein has a serine-rich region that is conserved among the Rec8 cohesins in plants. The serine-rich region is essential for the cohesin to interact with other proteins in the meiotic network. Furthermore, there are two potential proteolytic cleavage sites (PEST motifs) characterized as signals for rapid protein degradation in the predicted protein molecule. These results supported the candidate gene as a *Rec8-*like homologue in tetraploid wheat (*T. turgidum*), designated *TtRec8* (Figure S1).

### Expression profile of TtRec8 in LDN and LDN haploid

A significantly higher level of *TtRec8* transcripts was detected in the anthers at early meiotic prophase I than in roots and leaves of LDN by real-time PCR with the *TtRec8*-specific primer pair GM010F/GM010R (Table S2). The transcription level of *TtRec8* was the highest at interphase through early prophase I, and peaked at pachytene stage. After that, *TtRec8* transcripts continuously declined to a relative level of 3.77-4.14% at the end of meiosis I and later stages (Figure 1A). The relative transcription levels of *TtRec8* in roots and leaves were only about 9.04% and 0.02% of that in the anthers at interphase, respectively. *TtRec8* in haploid LDN exhibited an expression pattern similar to LDN, but its expression levels were relatively lower than LDN at the meiotic stages. The *TtRec8* transcript level in haploid LDN was about 60.61% of that in LDN at leptotene stage (Figure 1B). Overall, *TtRec8* showed an expression profile similar to the *Rec8*-like genes in model species (Lee *et al.*, 2005; Watanabe, 2005), further supporting *TtRec8* as a *Rec8*-like homologue in tetraploid wheat.

**Figure 1.**
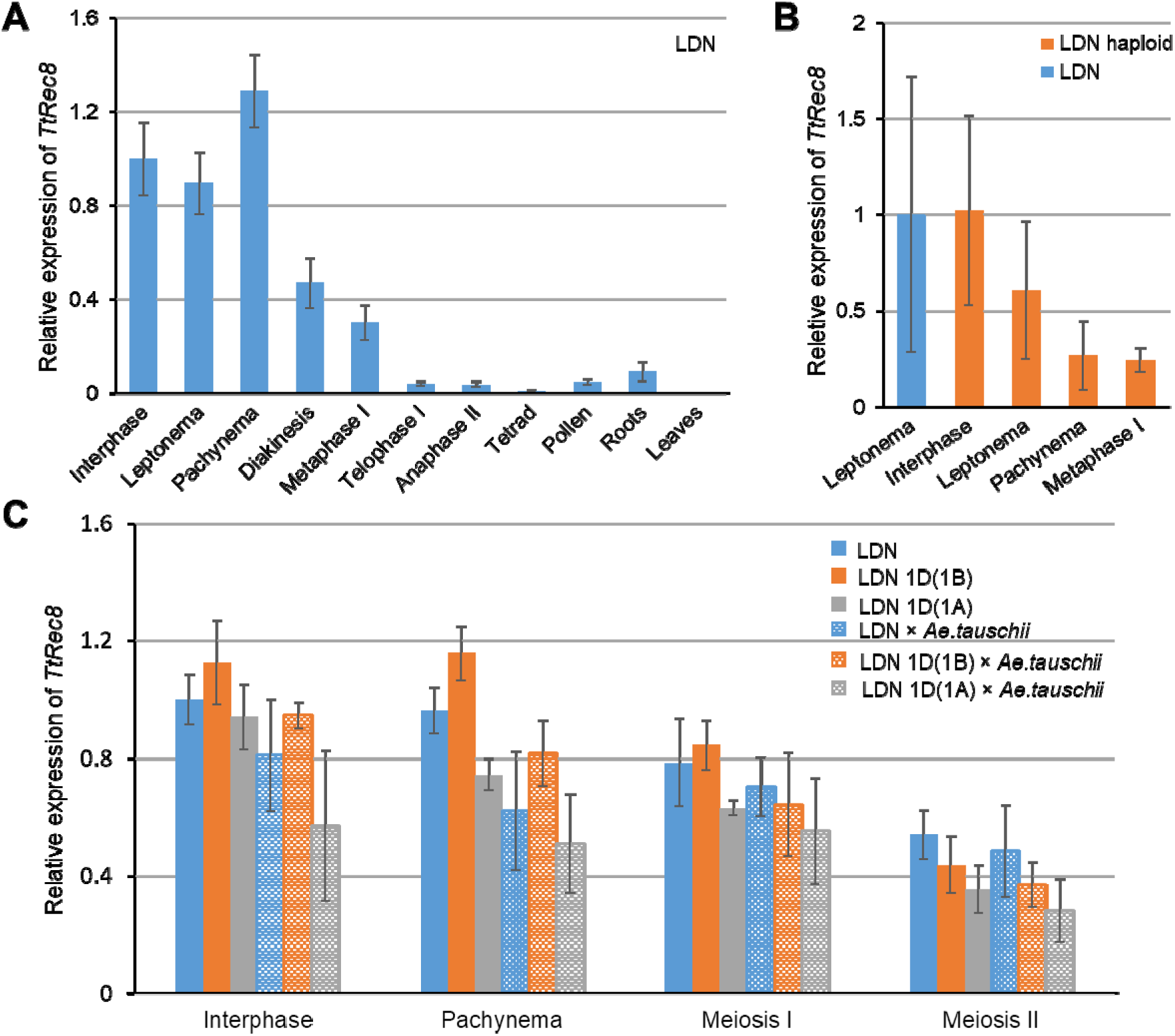
Relative transcript levels of *TtRec8* in roots, leaves, and anthers at different meiotic stages. **A)** LDN; **B)** LDN and LDN haploid; **C)** LDN, LDN 1D(1A), LDN 1D(1B), and their hybrids with *Ae. tauschii*. The comparative C_*T*_ method was used to determine the changes of *Rec8-*like gene expression in different samples (test) relative to the anthers at interphase (control). Fold difference in gene expression is 2^−ΔΔC*T*^, where ΔΔC_*T*_ = C_*T,test*_ – C_*T,control*_. Error bars represent standard deviation from the mean of three biological replicates.

The polyclonal antibody against TtRec8 was produced to probe TtRec8 protein in Western blotting and to localize TtRec8 on meiotic chromosomes. Immunoprecipitation was performed to verify the specificity of the anti-TtRec8 antibody. After anti-Rec8 antibody was incubated with the total protein extract from the anthers undergoing meiosis, a protein with a molecular weight a little over 60 kDa was immunoprecipitated. This molecular weight matched the predicted molecular weight 67.6 kDa of TtRec8. This protein was not present in the supernatant after immunoprecipitation. In addition, no precipitation was observed when anti-TtRec8 antibody was not added to the protein extract (Figure 2, *top*). These results validated the specificity of the antibody for TtRec8 protein in tetraploid wheat.

**Figure 2.**
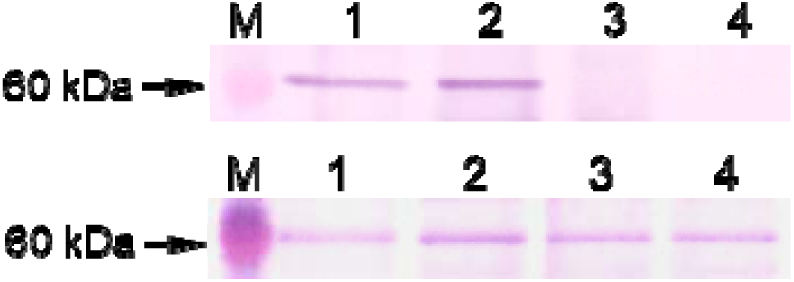
Immunoprecipitation and Western blotting of TtRec8 in LDN. *Top*: M-protein size marker; 1-total protein extracted from the anthers primarily at early prophase I; 2-proteins immunoprecipitated by anti-TtRec8 antibody; 3-negative control without TtRec8 and other proteins (Note: this sample was prepared from the immunoprecipitation experiment without anti-TtRec8 antibody added in the immunoprecipitation reaction); and 4-supernatant from the immunoprecipitation experiment with anti-TtRec8 antibody added in the reaction. ***Bottom*:** M-protein size marker; 1-interphase; 2-early prophase I; 3-metaphase I/anaphase I; and 4-metaphase II/anaphase II.

Western blotting with TtRec8 antibody detected the highest level of TtRec8 protein in the anthers at early prophase I. Moderate amounts of TtRec8 were detected in the anther samples at later meiotic stages, indicating partial retaining of TtRec8 cohesin in the meiocytes. The TtRec8 protein level in the anther sample at early prophase I was higher than those at later meiotic stages, but the difference did not seem to be significant (Figure 2, *bottom*). This might result from the meiotically unsynchronized meiocytes present in some of the anthers sampled for the Western blotting analysis. The anther samples we collected for each of the meiotic stages occasionally contained some off-type meiocytes, i.e. those at a meiotic stage earlier or later than the targeted stage. The off-type meiocytes could lower the TtRec8 protein level in the anther sample at early prophase I in which *TtRec8* has the highest expression. On the other hand, the anther samples at later meiotic stages occasionally contained the meiocytes at early prophase I, leading to an elevated TtRec8 protein level. Thus, the off-type meiocytes narrowed down the overall difference in the TtRec8 protein levels between the anther sample at early prophase I and those at the later meiotic stages.

### Subcellular localization of TtRec8 protein

The TtRec8 antibody was employed to localize endogenous TtRec8 protein on the meiotic chromosomes in the meiotically staged male meiocyte nuclei using the *in situ* immunolocalization technique. TtRec8 protein was clearly detected along the entire chromosomes from early leptonema through pachynema in meiosis I (Figure 3, a1-a3 and b1-b3). After pachynema, TtRec8 was almost undetectable by *in situ* immunolocalization (Figure 3, c1-j3). Apparently, the majority of the TtRec8 protein dissociated from chromosomes after pachynema. TtRec8 protein was not detected on mitotic chromosomes in the somatic cells of anthers that underwent mitosis, indicating TtRec8 is meiosis-specific (Figure 4). The kinetics of TtRec8 through the meiotic processes in LDN was very similar to the Rec8 cohesins in yeast and other model species. These are strong evidence validating the identity of TtRec8 as a Rec8-like homologue in tetraploid wheat.

**Figure 3.**
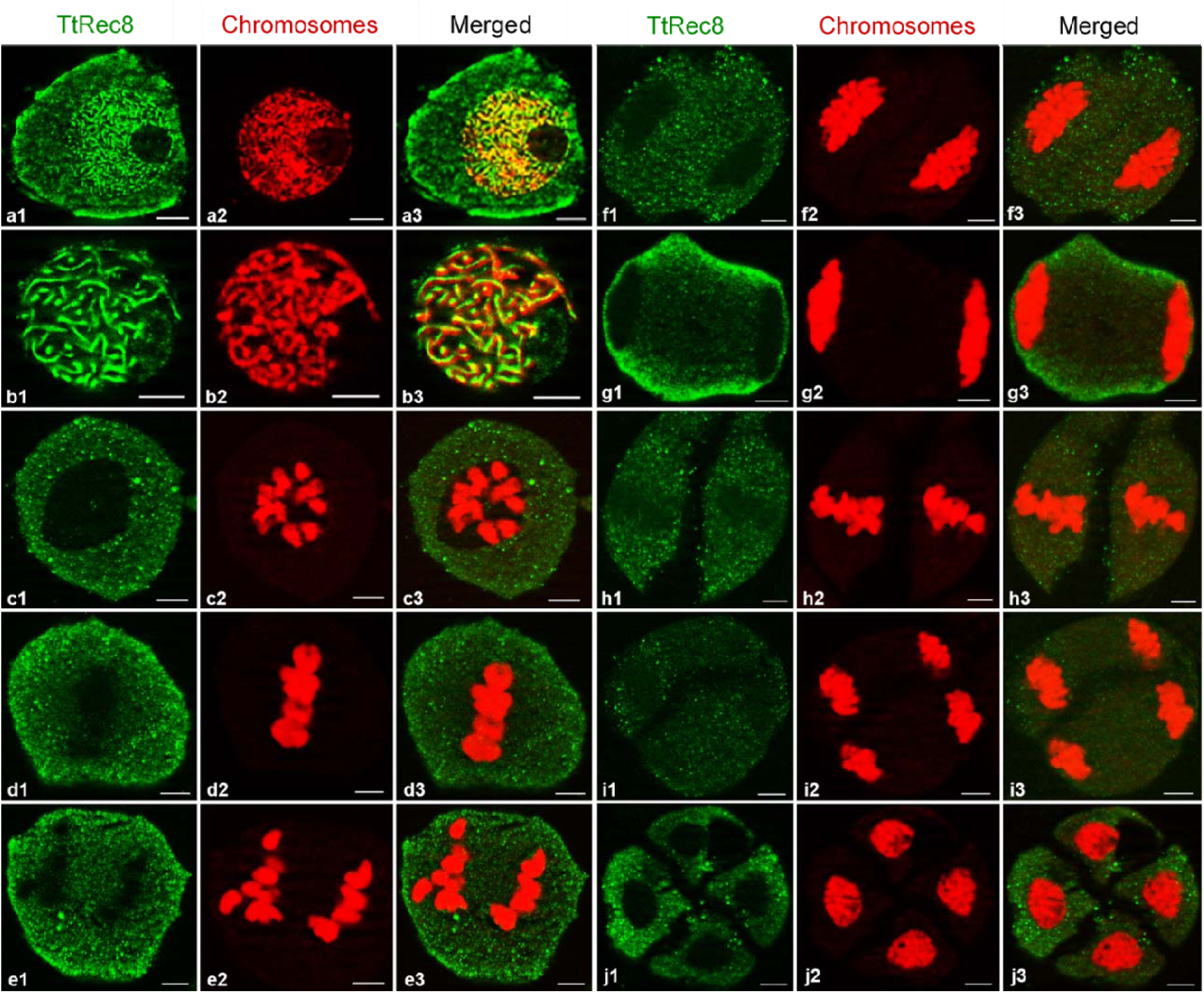
Fluorescent *in situ* immunolocalization of TtRec8 protein (green) on meiotic chromosomes (red) in LDN. **a1-a3:** leptonema; **b1-b3:** pachynema; **c1-c3:** diakinesis; **d1-d3:** metaphase I; **e1-e3:** anaphase I; **f1-f3:** early telophase I; **g1-g3:** late telophase I; **h1-h3:** metaphase II; **i1-i3:** anaphase II; and **j1-j3:** tetrad. Scale bar = 5 µm.

**Figure 4.**
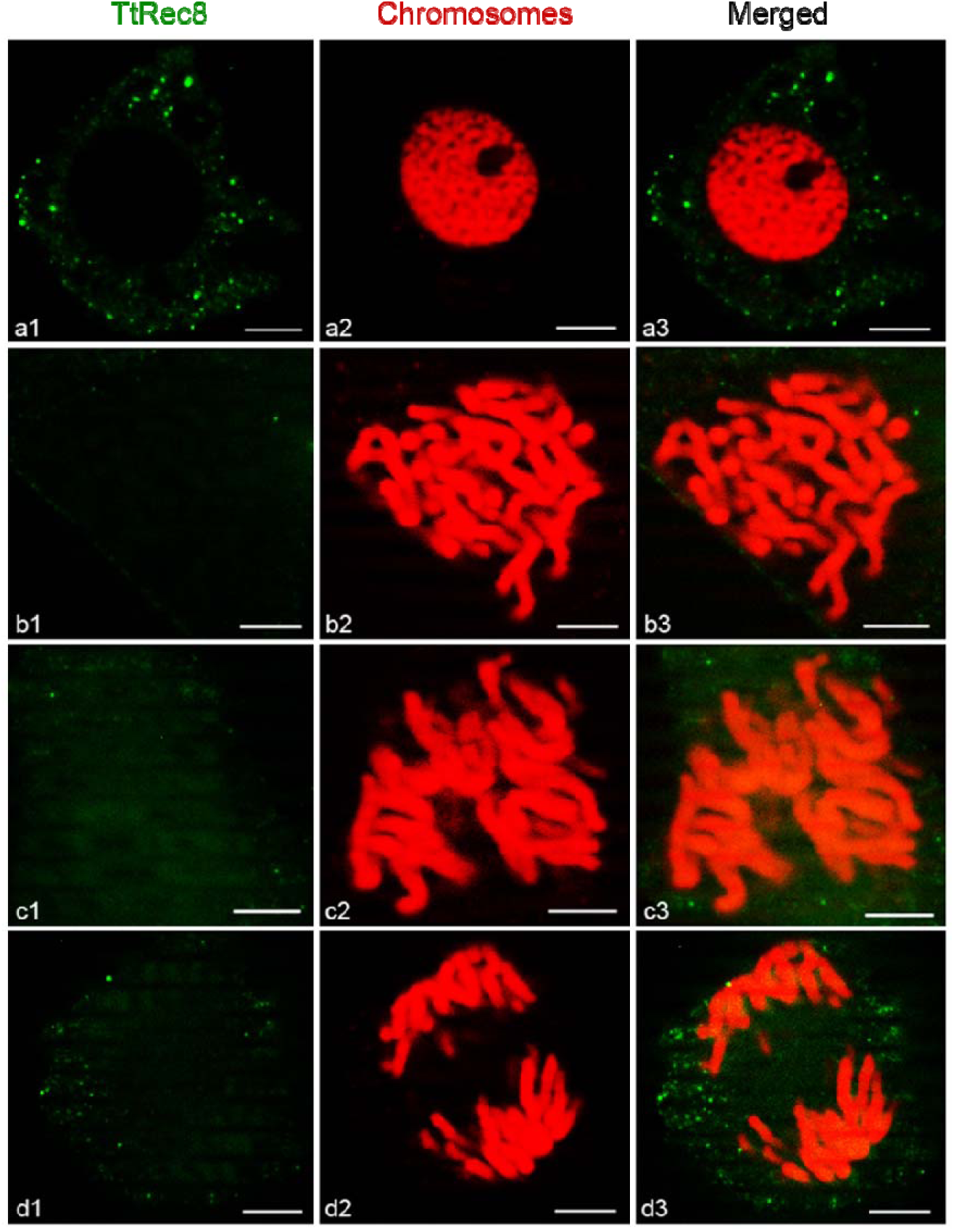
Fluorescent *in situ* immunolocalization of TtRec8 protein (green) on the mitotic chromosomes (red) of anthers in LDN. **a1-a3:** prophase; **b1-b3:** prometaphase; **c1-c3:** metaphase; and **d1-d3:** anaphase. Scale bar = 5 µm.

### Genomic sequence cloning and chromosomal localization of TtRec8

Screening of the LDN BAC library with the cDNA probes of *TtRec8* identified six BAC clones (No. 7, 18, 20, 24, 30, and 31) containing *TtRec8* (Figure S2A). Fingerprinting with *Hin*dIII and CAPS analysis categorized these six BAC clones into two groups, i.e. clones No. 7, 18, 24, and 30 in one group, and clones No. 20 and 31 in another group (Supplemental Figures 2B & 2C). Apparently, these two groups of the BAC clones each harbored a different homoeoallele of *TtRec8*. The two *TtRec8* homoeoalleles identified in LDN mapped to chromosome 1A and 1B using CS nulli-tetrasomic and LDN DS lines (Figure 5), and designated *TtRec8-A1* and *TtRec8-B1*, respectively.

**Figure 5.**
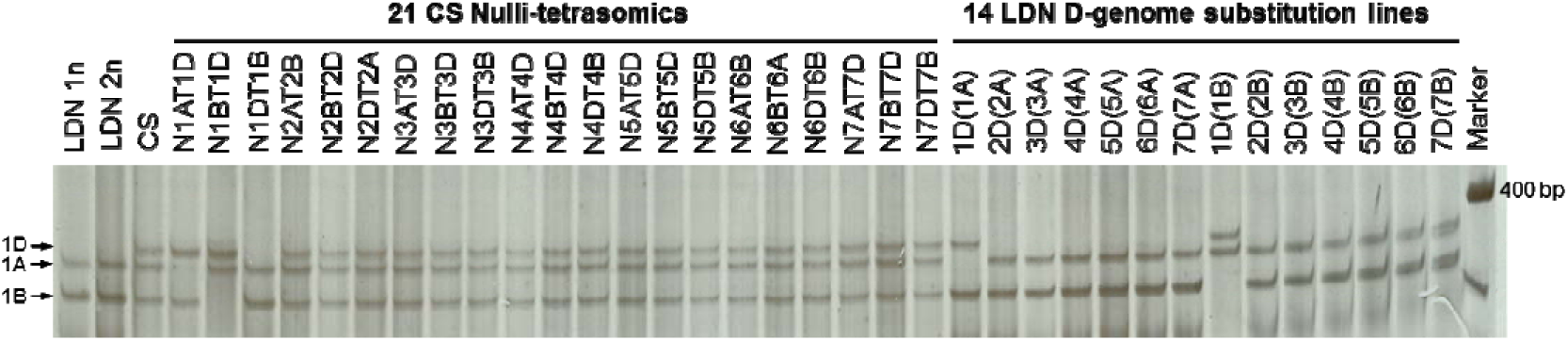
Chromosomal localization of the wheat *Rec8* homologues by *Dde*I-CAPS with *TtRec8*-specific primers (GM008F/GM008R). Two fragments were detected in tetraploid wheat LDN and three in hexaploid wheat CS, indicating the presence of two *TtRec8* homoeoalleles in tetraploid wheat and three in hexaploid wheat. They were positioned to chromosome 1D, 1A and 1B, respectively.

In addition, chromosome 1D of CS wheat (*T. aestivum*) was found to contain another homoeoallele of the *Rec8* gene (Figure 5), designated *TaRec8-D1*. The rice *Rec8*-like gene *OsRad21-4* (GenBank accession NP_001056426.1) and *Brachypodium* gene encoding sister chromatid cohesion 1 protein 1-like protein (GenBank accession XP_003567819.1) were assigned to the long arms of chromosome 5 and chromosome 2, respectively. Both chromosomes are collinear with wheat chromosomes in homoeologous group 1, *i.e.* 1A, 1B, and 1D (Zhang *et al.*, 2006; Kumar *et al.*, 2012). These results further confirm the identity of the wheat homoeoalleles as the *Rec8*-like genes in polyploid wheat.

A 40-kb genomic DNA fragment containing *TtRec8-A1* in the BAC clone No. 18 was sub-cloned into a cosmid vector for sequencing. A 6.5 kb DNA segment harboring *TtRec8-A1* was completely sequenced. A similar approach was used to isolate the genomic sequence of the homoeoallele *TtRec8-B1* from the BAC clone No. 20. A genomic region of 6.6 kb harboring *TtRec8-B1* was completely sequenced. DNA sequence analysis indicated that the initially cloned cDNA is for the homoeoallele *TtRec8-A1* located on chromosome 1A.

### Cloning of TtRec8-B1 and TaRec8-D1 cDNAs

We cloned cDNA of the homoeoallele *TtRec8-A1* located on chromosome 1A from LDN, but not *TtRec8-B1* on chromosome 1B in the initial cloning experiment. Thus, we attempted to clone cDNA of *TtRec8-B1* from the substitution line LDN 1D(1A), where LDN chromosome 1A was replaced by CS chromosome 1D. Meanwhile, the LDN substitution lines LDN 1D(1A) and LDN 1D(1B), where CS chromosome 1D respectively replaced LDN chromosome 1A and 1B, were used to clone cDNA of the homoeoallele *TaRec8-D1* located on CS chromosome 1D. We found that the cDNA primer pair GM067F/GM065R spans the start and stop codons of all three homoeoalleles (*TtRec8-A1, TtRec8-B1*, and *TaRec8-D1*) according to their genomic DNA and *TtRec8-A1* cDNA sequences (Table S2). Therefore, GM067F/GM065R was used to amplify cDNAs of the three homoeoalleles from the cDNA pools prepared from the meiotic anthers at early prophase I of LDN 1D(1B) and LDN 1D(1A), respectively. The amplicons were cloned and sequenced. Sequence analysis indicated that the amplicons obtained from LDN 1D(1B) were the cDNA mixture of *TtRec8-A1* and *TaRec8-D1*, and amplicons from LDN 1D(1A) were the cDNA mixture of *TtRec8-B1* and *TaRec8-D1*. Comparative analysis identified the full-length cDNAs of *TaRec8-D1* and *TtRec8-B1*. They have the same length of 1,827 bp encoding 608 amino acids as *TtRec8-A1*. The cDNAs of *TtRec8-A1, TtRec8-B1*, and *TaRec8-D1* differed from each other at 52 SNP loci (Figure S3).

### Gene structures and predicted protein sequences of the Rec8 homoeoalleles

Alignment of the genomic DNA sequences with the coding DNA sequences of *TtRec8* indicates that both *TtRec8-A1* and *TtRec8-B1* contain 20 exons and 19 introns. The largest exon has 268 bp and the smallest 20 bp in length. The largest intron has 1,491 bp (between exon 6 and 7) and the smallest is 71 bp (between exon 15 and 16) in length. Interestingly, we found that *TtRec8* and the rice *Rec8*-like gene *OsRad21-4* share extremely high similarities in the number, size, and distribution of exons/introns despite a slight difference in length of the genomic DNA sequences (Figure 6A). Thus, the *Rec8*-like gene is highly conserved in rice and wheat. The genomic DNA sequence of LDN *TtRec8-A1* showed high homology with a CS genomic DNA segment assigned to the long arm of chromosome 1A in the IWGSC CS RefSeq v1.0 assembly (TGACv1_scaffold_000516_1AL; https://wheat-urgi.versailles.inra.fr/Seq-Repository/Assemblies) (99.9% nucleotide sequence similarity in 6.5 kb region), indicating the location of *TtRec8* on the long arm of the group 1 chromosomes (i.e. chromosomes 1A and 1B).

**Figure 6.**
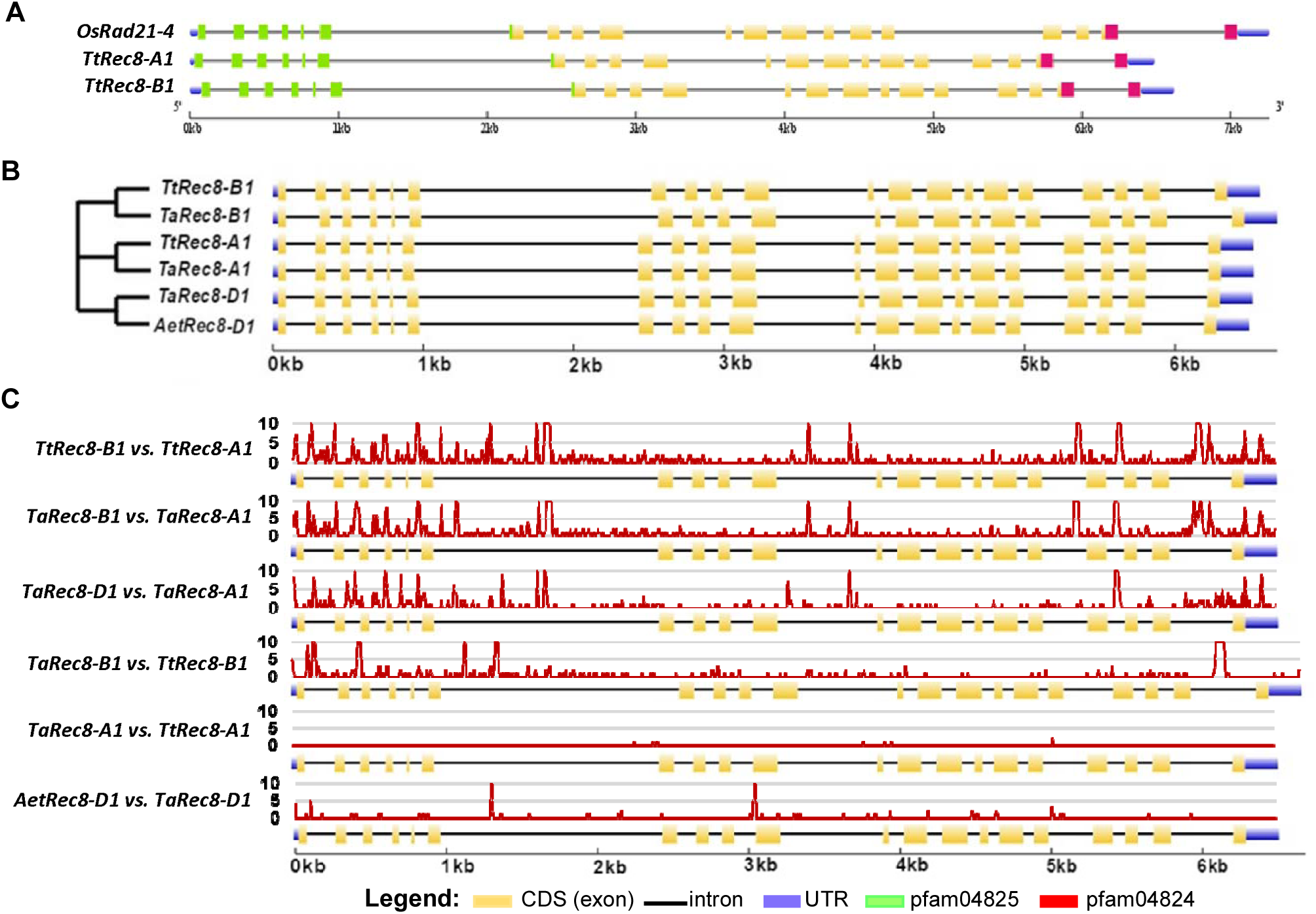
The gene structure and comparative analysis of the *Rec8* homoeoalleles in wheat and its D-genome ancestor and the *Rec8* orthologue in rice (*OsRad21-4*). **A)** The gene structures and comparative analysis of the *Rec8* orthologues in tetraploid wheat and rice. **B)** The gene structure (*right*) and phylogenetic tree (*left*) of the *Rec8* homoeoalleles in wheat and its D-genome ancestor. **C)** Pairwise comparative analysis of the *Rec8* homoeoalleles in wheat and its D-genome ancestor. The DNA sequence variation curves (red) were generated on a sliding window of 10 bp. The Y axes indicate the numbers of polymorphic nucleotides per 10 bp on the alignment.

In addition, we characterized the *Rec8* homoeoalleles on chromosomes 1A, 1B, and 1D of bread wheat (*T. aestivum*) (*TaRec8-A1, TaRec8-B1*, and *TaRec8-D1*) and the one on chromosome 1D of *Ae. tauschii* (*AetRec8-D1*). Genomic DNA sequences of *TaRec8-A1* and *TaRec8-B1* were extracted from the IWGSC CS RefSeq v1.0 assembly (https://wheat-urgi.versailles.inra.fr/Seq-Repository/Assemblies) by BLASTn with the genomic DNA sequences of *TtRec8-A1* and *TtRec8-B1* as queries. Partial cDNA sequences of *TaRec8-A1* and *TaRec8-B1* (∼1.6 kb at 3’ end) were identified from the Hexaploid Wheat Transcriptome Database (https://wheat.pw.usda.gov/WheatExp/; Pearce *et al.*, 2015) by BLASTn with the cDNA sequences of *TtRec8-A1* and *TtRec8-B1* as queries. Remaining 5’ cDNA sequences for *TaRec8-A1* (225 bp) and *TaRec8-B1* (192 bp) were deduced according to the splicing patterns of the corresponding regions in *TtRec8-A1* and *TtRec8-B1*. The genomic DNA sequence of *TaRec8-D1* was extracted from the IWGSC CS RefSeq v1.0 assembly by BLASTn with *TaRec8-D1* cDNA sequence as query. The genomic DNA sequence of *AetRec8-D1* was obtained from the *Ae. tauschii* reference genome sequence (http://aegilops.wheat.ucdavis.edu/ATGSP/data.php) by BLASTn with the genomic DNA sequence of *TaRec8-D1* as query. *AetRec8-D1* shares over 99% similarity with *TaRec8-D1* in the genomic DNA sequence. Thus, the cDNA sequence of *TaRec8-D1* was used in the gene structural analysis for *AetRec8-D1*.

Phylogenetic analysis placed the *Rec8* homoeoalleles on chromosomes 1A, 1B, and 1D of wheat and the D-genome ancestor *Ae. tauschii* (*AetRec8-D1*) into three distinct clusters (Figure 6B). Significant differences were observed between the homoeoalleles from different subgenomes/chromosomes, i.e. *TtRec8-B1* vs. *TtRec8-A1, TaRec8-B1* vs. *TaRec8-A1*, and *TaRec8-D1* vs. *TaRec8-A1* (Figure 6C). However, we found that the cDNA sequences of *TaRec8-A1* and *TtRec8-A1* were identical even though their genomic DNA sequences differed at single nucleotide positions in several introns. *TaRec8-D1* was slightly different from *AetRec8-D1* in the DNA sequences. In contrast, *TaRec8-B1* was quite different from *TtRec8-B1* in the DNA sequences of both introns and exons. Overall, the *Rec8* homoeoalleles from different subgenomes have a greater variation in DNA sequences than the homoalleles within a subgenome in wheat.

The amino acid sequences of the proteins encoded by *TtRec8-B1, TtRec8-A1*, and *TaRec8-D1* were predicted based on their cDNA sequences cloned in this study. All three predicted proteins contain the conserved domains of the Rec8 cohesin (pfam04825 and pfam04824), the serine-rich region conserved in plant cohesins, and the potential PEST motifs present in the cohesion proteins (Figure S4). Alignment of their amino acid sequences identified a total of 23 amino acid positions polymorphic among these three homoeologous Rec8 proteins. Out of these 23 amino acid positions, 17 were polymorphic between TtRec8-B1 and TtRec8-A1/TaRec8-D1 and 6 polymorphic between TtRec8-A1 and TtRec8-B1/TaRec8-D1 (Figure S4). These amino acid polymorphisms structurally differentiate these three Rec8 cohesion proteins from each other, and might lead to functional differentiation of the homoeologous proteins as meiotic cohesin in polyploid wheat.

A phylogenetic tree was constructed from the predicted protein sequences of the wheat *Rec8* homoeoalleles and their orthologues in other plants, indicating that TtRec8-A1 and TaRec8-D1 were more closely related to each other than their relationship with TtRec8-B1. The Rec8 cohesin-like proteins from barely and Brachypodium were clustered together with the three wheat Rec8 homoeoalleles, demonstrating a close relationship of the barley and Brachypodium Rec8-like proteins with those in polyploid wheat. The Rec8 cohesion proteins from maize, sorghum, foxtail millet, Japonica-type rice, and wild rice were clustered into another group related to the wheat-barley-Brachypodium cluster. The Syn1 protein from the dicotyledon Arabidopsis was further distinct from the cohesin proteins in the monocotyledons (Figure 7).

**Figure 7.**
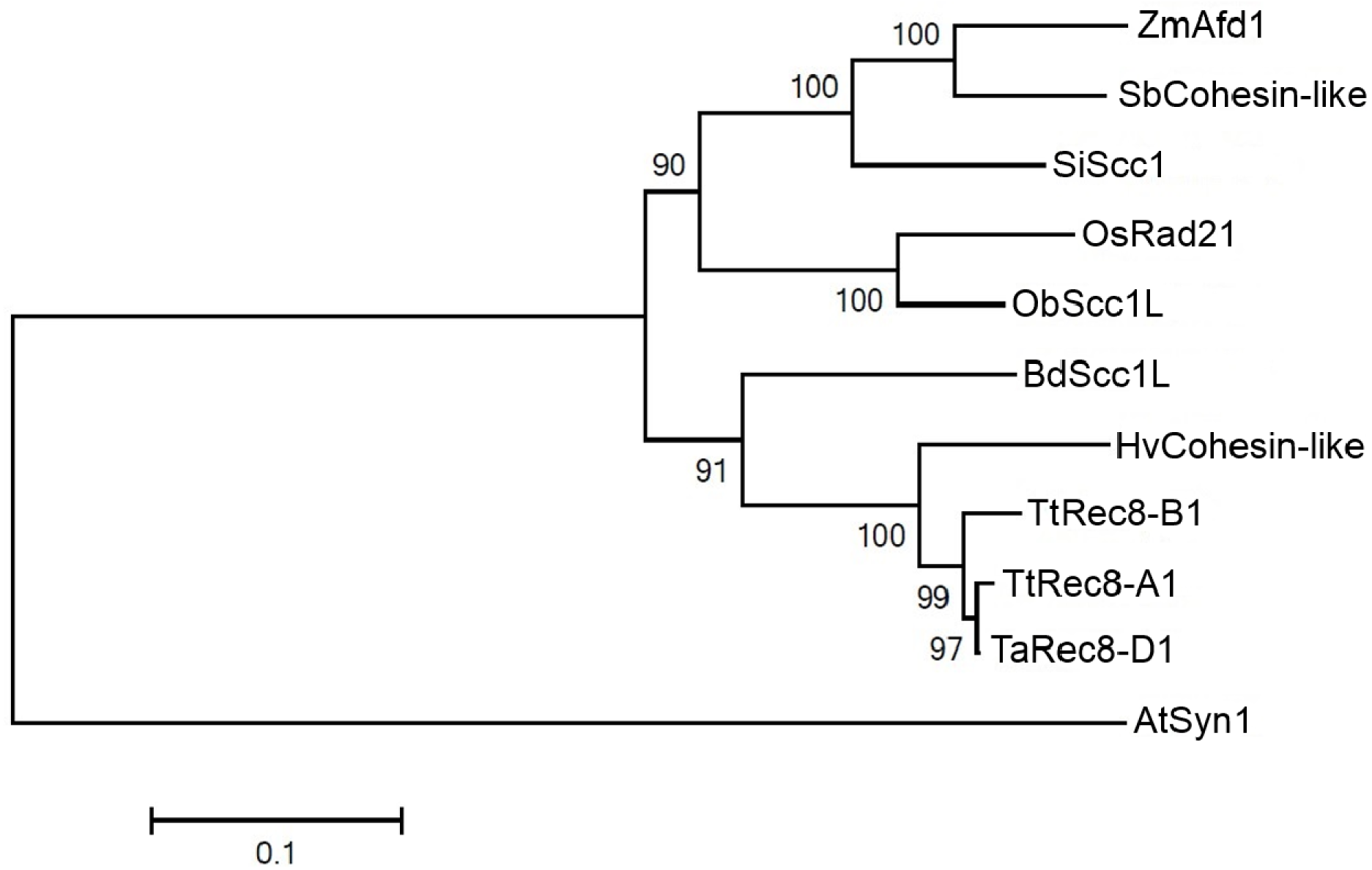
Phylogenetic tree (bootstrap value =1,000) of the Rec8 orthologues in wheat and other plants. It consists of TtRec8-A1, TtRec8-B1, TaRec8-D1, *Sorghum bicolor* SbCohesin-like (GenBank Accession XP_002440318.1), *Hordeum vulgare* HvCohesin-like (GenBank Accession BAJ93700.1), *Brachypodium distachyon* BdScc1L (GenBank Accession XP_003567819.1), *Setaria italica* SiScc1 (GenBank Accession XP_012699922.1), *Zea mays* ZmAfd1 (GenBank Accession NP_001105829.1), *Oryza sativa* OsRad21 (GenBank Accession NP_001056426.1), *Oryza brachyantha* ObScc1L (GenBank Accession XP_006654842.1), and *Arabidopsis thaliana* AtSyn1 (GenBank Accession NP_196168.1).

### Differential expression analysis of the Rec8 homoeoalleles in polyploid wheat

The real-time PCR with the cDNA primers (GM010F/GM010R) shared by the three wheat *Rec8* homoeoalleles (*TtRec8-A1, TtRec8-B1*, and *TaRec8-D1*) (Figure 8 and Figure S3) revealed the highest transcription level in LDN 1D(1B) (*TtRec8-A1* + *TaRec8-D1*), followed by LDN (*TtRec8-A1* + *TtRec8-B1*) and LDN 1D(1A) (*TtRec8-B1* + *TaRec8-D1*). *TtRec8-A1* showed the highest transcription level among these three *Rec8* homoeoalleles in tetraploid wheat (Figure 1C). Similar transcription profiles were observed with the *Rec8* homoeoalleles in the hybrids of LDN, LDN 1D(1A), and LDN 1D(1B) with *Ae. tauschii* (Figure 1C). Therefore, the homoeoalleles *TtRec8-A1, TtRec8-B1*, and *TaRec8-D1* expressed differentially in the tetraploid wheat background.

**Figure 8.**
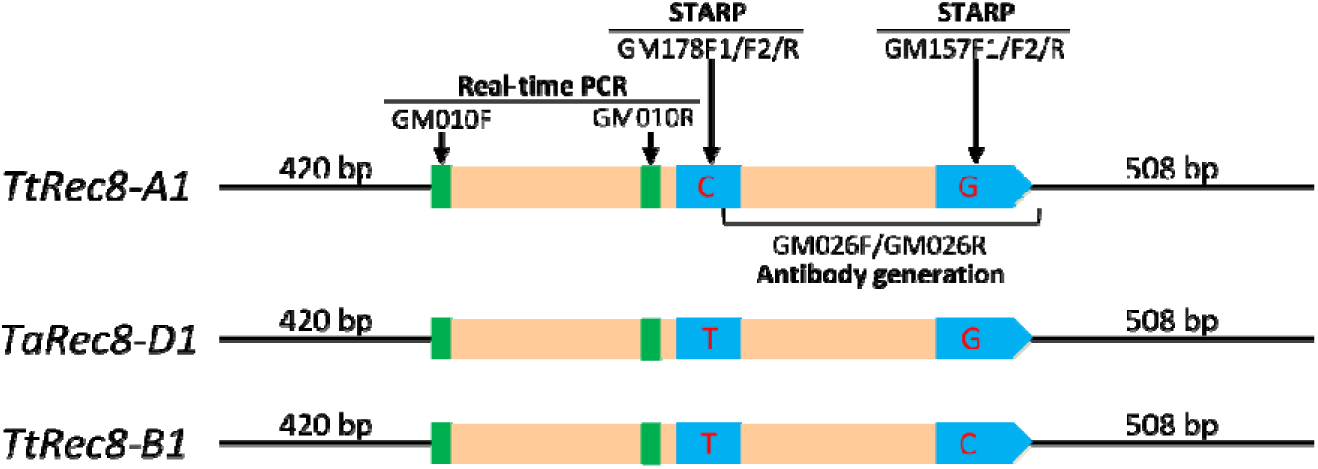
Graphical presentation of the cDNAs for the wheat *Rec8* homoeoalleles, showing the regions amplified by real-time PCR and STARP and the region used for TtRec8 antibody generation. Green bars refer to the primer positions of the real-time PCR, and blue bars refer to the regions spanned by STARP. Letters within the blue bars refer to the SNPs targeted by STARP.

The CDS of the three wheat *Rec8* homoeoalleles have extremely high similarity (97% between *TtRec8-A1* and *TtRec8-B1*, 98% between *TtRec8-B1* and *TaRec8-D1*, and 99% between *TtRec8-A1* and *TaRec8-D1*). They differed from each other only at single nucleotide positions. No indel was detected in their CDS (Figure S3). To further characterize the expression of these three *Rec8* homoeoalleles in polyploid wheat, we examined their relative levels of transcription using the new SNP-based STARP technique (Long *et al.*, 2017). A group of three allele-specific (AS) primers, including two allele-specific forward primers and one common reverse primer for each of the two SNP loci targeted (GM157F1/GM157F2/GM157R and GM178F1/GM178F2/GM178R), were designed according to the contextual sequences of the SNPs (Figure 8; Figure S3; Table S2). Each of the SNPs is located within an exon of the homoeoalleles, with no intron in the region spanned by the STARP primer sets. Thus, the same amplicons were expected for each of the primer sets in the genomic DNA and cDNA of the homoeoalleles. Prior to the transcript amplification, the STARP primer sets were tested for primer specificity and amplification efficiency in the genomic DNA of tetraploid wheat LDN, the substitution lines LDN 1D(1A) and LDN 1D(1B), *T. urartu* (wheat A genome donor), and *Ae. tauschii* (wheat D genome donor). As shown in Figure 9A (*top*), GM157F1 is specific for *TtRec8-A1, TaRec8-D1, TuRec8-A1*, and *AetRec8-D1*, while GM157F2 is specific for *TtRec8-B1*. For the primer set GM178F1/F2/R, GM178F1 is specific for *TtRec8-A1* and *TuRec8-A1*, while GM178F2 is specific for *TtRec8-B1, TaRec8-D1*, and *AetRec8-D1* (Figure 9A, *bottom*). These two STARP primer sets amplified a transcript similar to that of *TtRec8-A1* in *T. urartu* (*TuRec8-A1*) and the one similar to that of *TaRec8-D1* in *Ae. tauschii* (*AetRec8-D1*) (Figure 9A). The pairwise amplification tests indicated that the STARP primer sets equally amplified the homoeoallele pairs in the genomic DNAs of LDN, LDN 1D(1A), and LDN 1D(1B) at different PCR cycles (Figure 9B). Therefore, these two STARP primer sets were suitable for differentially examining the transcripts of the individual *Rec8* homoeoalleles in LDN wheat background.

**Figure 9.**
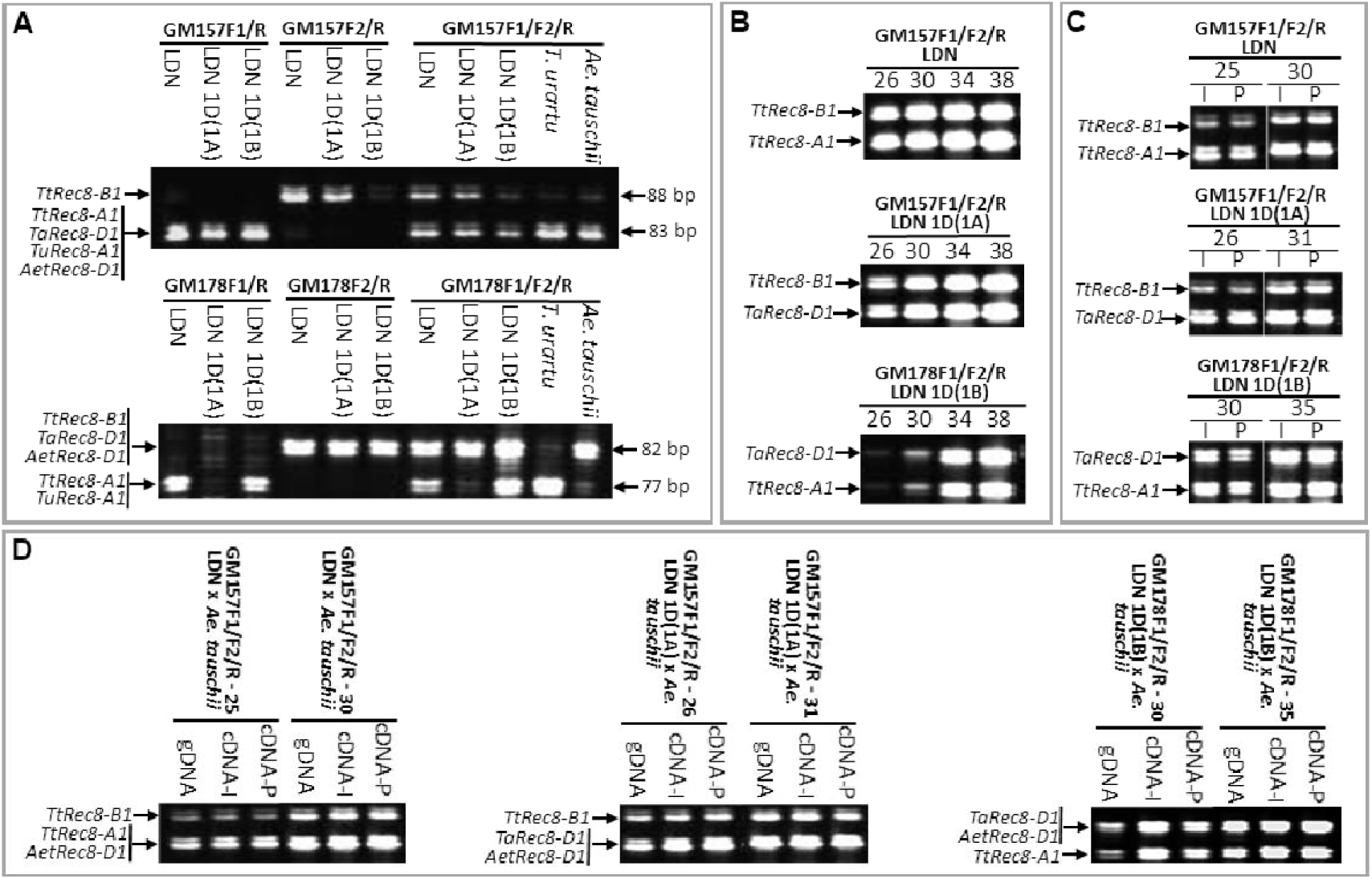
STARP-based expression analysis of the *Rec8* homoeoalleles (*TtRec8-A1, TtRec8-B1, TaRec8-D1, TuRec8-A1*, and *AetRec8-D1*) in polyploid wheat. **A)** Primer specificity tests (32 PCR cycles) in the genomic DNA of LDN, LDN 1D(1A), LDN 1D(1B), *T. urartu* (wheat A genome donor), and *Ae. tauschii* (wheat D genome donor). **B)** PCR amplification efficiency tests of the allele-specific primers in the genomic DNA of LDN, LDN 1D(1A), and LDN 1D(1B) at different PCR cycles (26, 30, 34, and 38). **C)** PCR amplification of the cDNA from the anthers at the meiotic interphase (I) and pachytene (P) stages in LDN, LDN 1D(1A), and LDN 1D(1B) with different primer sets at different PCR cycles (25, 30; 26, 31; and 30, 35). **D)** PCR amplification of the genomic DNA and cDNA from the anthers at I and P stages of LDN × *Ae. tauschii*, LDN 1D(1A) × *Ae. tauschii*, and LDN 1D(1B) × *Ae. tauschii* hybrids with different primer sets at different PCR cycles (25, 30; 26, 31; and 30, 35).

The primer set GM157F1/F2/R discriminated *TtRec8-B1* from *TtRec8-A1* and *TaRec8-D1* in LDN and LDN 1D(1A), respectively (Figure 9C, *top* & *middle*), while GM178F1/F2/R discriminated *TtRec8-A1* from *TaRec8-D1* in LDN 1D(1B) (Figure 9C, *bottom*). Both *TtRec8-A1* and *TaRec8-D1* showed significantly higher expression levels than *TtRec8-B1* in the anthers at interphase (I) and pachytene (P) stages (Figure 9C, *top* & *middle*). The expression levels of *TtRec8-A1* and *TaRec8-D1* seemed to be comparable at both meiotic stages (Figure 9C, *bottom*). Therefore, *TtRec8-A1* should be the primary gene for Rec8 cohesin production in tetraploid wheat. *TtRec8-B1* might also contribute to Rec8 cohesin production, but should not be significant. When chromosome 1A of tetraploid wheat LDN was replaced by chromosome 1D of hexaploid wheat CS, *TaRec8-D1* expressed at a significantly higher level than *TtRec8-B1* (Figure 9C, *middle*). Apparently, *TtRec8-B1* intrinsically expressed at a lower level than other *Rec8* homoeoalleles of wheat.

In addition, we examined expression of the *Rec8* homoeoalleles in the hybrids of LDN, LDN 1D(1A), and LDN 1D(1B) with *Ae. tauschii*. *TtRec8-B1* still showed the lowest expression level among the *Rec8* homoeoalleles in the hybrids as observed in their tetraploid wheat parents (Figures 9C & 9D). The presence of the *Ae. tauschii* D genome harboring *AetRec8-D1* in the hybrids did not seem to increase overall expression of the chromosome 1D-specific *Rec8* alleles (Figure 9D). Therefore, the *Rec8* homoeoalleles on chromosome 1A and 1D were transcriptionally more active than the one on chromosome 1B in tetraploid wheat and its hybrids with *Ae. tauschii*, and probably in hexaploid wheat as well.

To further characterize expression of the wheat *Rec8* homoeoalleles, we examined Rec8 cohesin on meiotic chromosomes in the specially constructed genotypes with different homoeoallelic combinations of the wheat *Rec8* gene by *in situ* immunolocalization. Rec8 cohesin appeared to be evenly distributed along the individual chromosomes at early prophase I in the LDN × *Ae. tauschii* hybrid (*TtRec8-A1* + *TtRec8-B1* + *AetRec8-D1*), LDN haploid (*TtRec8-A1* + *TtRec8-B1*), LDN 1D(1A) (*TtRec8-B1* + *TaRec8-D1*), LDN 1D(1A) × *Ae. tauschii* hybrid (*TtRec8-B1* + *TaRec8-D1* + *AetRec8-D1*), LDN 1D(1B) (*TtRec8-A1* + *TaRec8-D1*), and LDN 1D(1B) × *Ae. tauschii* hybrid (*TtRec8-A1* + *TaRec8-D1* + *AetRec8-D1*) (Figure 10). We did not observe significant quantitative variation of unbound Rec8 protein in the cytoplasm/nucleus among the genotypes. Each of these genotypes contained one or more actively-transcribed *Rec8* homoeoalleles (*TtRec8-A1, TaRec8-D1*, and *AetRec8-D1*). The presence of extra actively- transcribed *Rec8* homoeoalleles did not noticeably increase the amount of the Rec8 cohesin on the individual chromosomes (Figures 10C and 10E). The genotype or mutant that contains only one of the wild homoeoalleles (*TtRec8-A1, TtRec8-B1, TaRec8-A1, TaRec8-B1*, and *TaRec8-D1*) is not available in wheat. Thus, we were unable to precisely assess the role of a single homoeoallele in Rec8 cohesin production in wheat. As indicated in the differential transcriptional analysis of the *Rec8* homoeoalleles, the contribution of the *Rec8* homoeoallele on chromosome 1B to Rec8 cohesin production might be minimal, if there is any, in polyploid wheat. The actively-transcribed *Rec8* homoeoalleles on chromosomes 1A and 1D may be the primary genes for Rec8 cohesin production in polyploid wheat.

**Figure 10.**
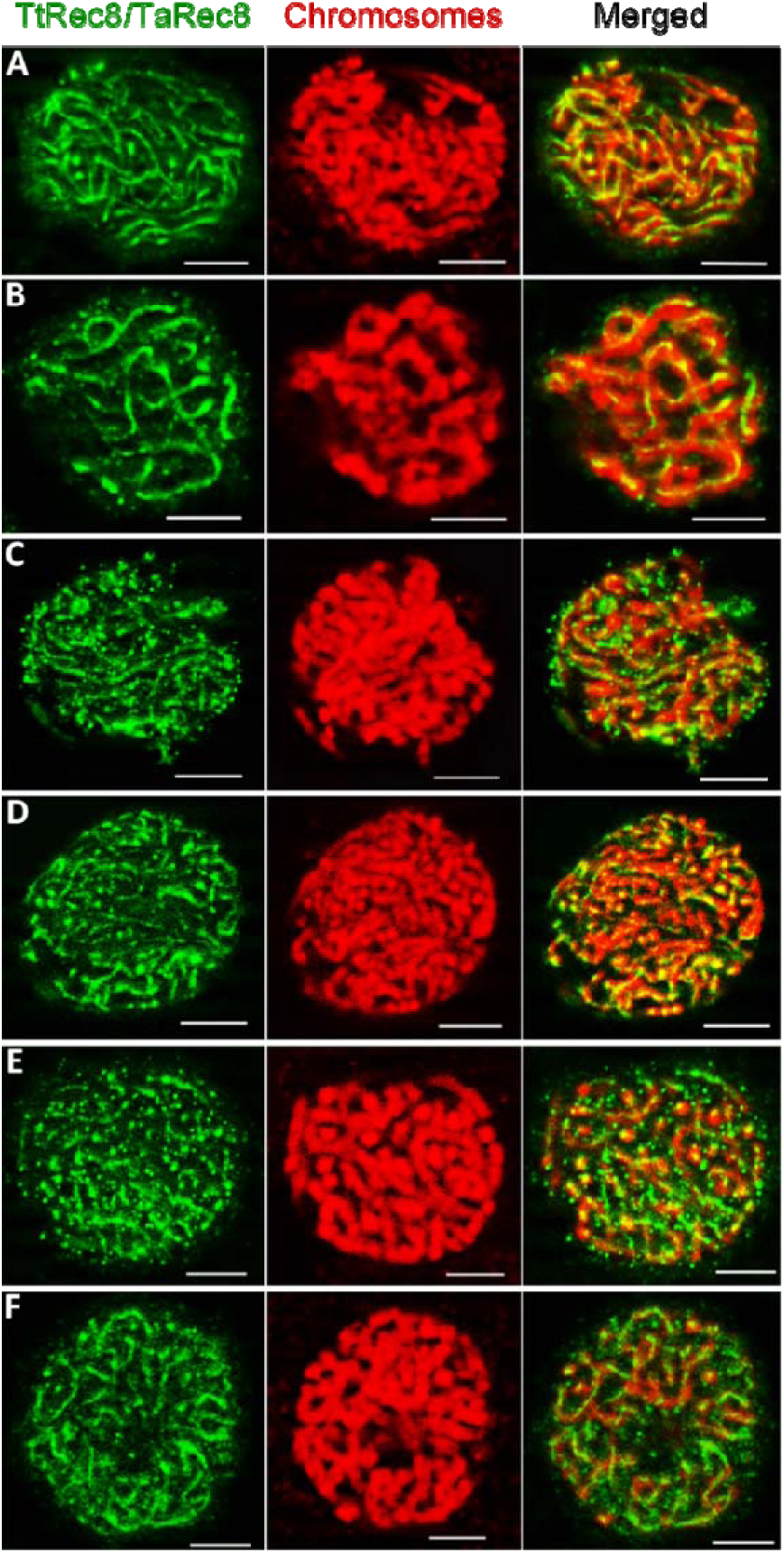
Fluorescent immunolocalization of TtRec8 protein (green) on the meiotic chromosomes (red) at early prophase I (leptonema-pachynema). **A)** LDN × *Ae. tauschii*; **B)** LDN haploid; **C)** LDN 1D(1A); **D)** LDN 1D(1A) × *Ae. tauschii*; **E)** LDN 1D(1B); and **F)** LDN 1D(1B) × *Ae. tauschii*. Scale bar = 5 µm.

## DISCUSSION

The Rec8-like cohesin functions as a meiotic cohesion protein in a very conserved manner from lower to higher eukaryotes (Watanabe and Nurse, 1999; Yu and Dawe, 2000; Chelysheva *et al.*, 2005). We took this advantage to clone the *Rec8*-like genes in polyploid wheat through a comparative genomics approach. This work could not be accomplished by map-based cloning, the most widely-used gene cloning method, due to the unavailability of an alternative allele or mutant for the *Rec8*-like genes in wheat. Knockout or mutation of the *Rec8*-like genes leads to abnormal meiosis and sterility (Bai *et al.*, 1999; Bhatt *et al.*, 1999; Cai *et al.*, 2003; Golubovskaya *et al.*, 2006; Zhang *et al.*, 2006). In addition, the complexity of the meiotic genetic network and difficulties in phenotyping individual meiotic events would cause extra encumbrance for map-based cloning of the meiotic genes. In the present study, we successfully cloned the *Rec8*-like homologues in tetraploid wheat by referencing the *Rec8*-like genes in model species, indicating the efficacy of this comparative genomics approach in cloning conserved meiotic genes especially from the species with a large and complex genome.

TtRec8 cohesin was detected on the meiotic chromosomes of wheat till pachynema at prophase I, but not after that by *in situ* immunolocalization. However, we detected lowered levels of TtRec8 protein in the anthers after early prophase I by Western blotting, indicating partial retaining of TtRec8 protein at later meiotic stages. Most likely, the retained TtRec8 protein was associated with the centromeric regions of meiotic chromosomes at later stages as observed in model species (Lee *et al.*, 2005; Watanabe, 2005). But, the retained TtRec8 cohesin was not imperceptible by *in situ* immunolocalization due probably to the condensed chromosomal structure after early prophase I. Similar observations were reported for the Rec8-like cohesins in Arabidopsis and maize (Cai *et al.*, 2003; Chelysheva *et al.*, 2005; Yuan *et al.*, 2012).

Wheat is an allopolyploid with two (tetraploid wheat, genome AABB) or three (hexaploid wheat, genome AABBDD) homoeologous subgenomes. A gene generally has two homoeoalleles in tetraploid wheat and three in hexaploid wheat (Murai *et al.*, 1999; Huang *et al.*, 2002; Kimbara *et al.*, 2004; Zhang *et al.*, 2011; Brenchley *et al.*, 2012; Leach *et al.*, 2014). The homoeoalleles of a gene in polyploid wheat generally share high levels of similarity in DNA sequence and function, making gene cloning and functional analysis a challenging task (Zhang *et al.*, 2011; Brenchley *et al.*, 2012). In this study, we recovered and cloned the cDNA of the homoeoallele *TtRec8-A1*, but not *TtRec8-B1* from tetraploid wheat in the initial RACE-based cloning experiments. This might result from the lower expression level of *TtRec8-B1* than *TtRec8-A1* in tetraploid wheat. The use of LDN 1D(1A) substitution line, where chromosome 1A containing *TtRec8-A1* was replaced by chromosome 1D, enhanced the abundance of *TtRec8-B1* transcripts in the cloning pool. In addition, this substitution line introduced the homoeoallele *TaRec8-D1* on chromosome 1D of hexaploid wheat into the transcript pool for cloning. Both *TtRec8-B1* and *TaRec8-D1* were successfully recovered and cloned from this engineered transcript pool. Apparently, this is an effective approach to clone a gene with low transcript abundance, especially in the polyploids with multiple similar homoeoalleles for the gene.

The *Rec8*-like homoeoalleles in the A, B, and D subgenomes of polyploid wheat are highly conserved in CDS and splicing patterns. However, there are substantial differences in their intronic regions. Interestingly, we found that the *Rec8* homoalleles in the wheat subgenome B (i.e. *TtRec8-B1* and *TaRec8-B1*) had a higher DNA sequence variation than those in the subgenomes A (i.e. *TtRec8-A1* and *TaRec8-A1*) and D (i.e. *TaRec8-D1* and *AetRec8-D1*), especially in the intronic regions. Wheat genome mapping also identified a similar differentiation of these three subgenomes in genetic diversity (Chao *et al.*, 1989; Felsenburg *et al.*, 1991; Siedler *et al.*, 1994; Petersen *et al.*, 2006). The distinction of the wheat B subgenome from the A and D subgenomes in genetic diversity may imply a different origin and evolutionary route for the B subgenome.

The three *Rec8* homoeoalleles we cloned in this study (*TtRec8-A1, TtRec8-B1*, and *TaRec8-D1*) differ from each other only at single nucleotide positions in CDS. It was a big challenge to perform differential expression analysis of the highly similar homoeoalleles under a polyploid condition. To confront the challenge, we partitioned the *Rec8* homoeoalleles in pairs using the cytogenetic stocks LDN 1D(1A) (*TtRec8-B1* and *TaRec8-D1*) and LDN 1D(1B) (*TtRec8-A1* and *TaRec8-D1*), and then differentially examined the relative expression levels of the individual pair of the *Rec8* homoeoalleles using the newly developed SNP-based PCR technique STARP (Long *et al.*, 2017). This integrative cytogenetic and genomic approach enabled us to perform pairwise differential expression analysis of the highly similar *Rec8* homoeoalleles under a polyploid condition. We found that both *TtRec8-A1* and *TaRec8-D1* expressed comparably at a significantly higher level than *TtRec8-B1* in tetraploid wheat and its hybrids with *Ae. tauschii* as well. Apparently, the *Rec8* homoeoalleles on chromosomes 1A and 1D expressed dominantly over the one on chromosome 1B in tetraploid wheat, and probably in hexaploid wheat as well. Concurrent presence of two or three actively-transcribed homoeoalleles (*TtRec8-A1, TaRec8-D1*, and *AetRec8-D1*) in LDN 1D(1B) and the hybrids with *Ae. tauschii* did not lead to a noticeable additive effect on Rec8 cohesin production according to the *in situ* immunolocalization results. It seems that either of the two actively-transcribed homoeoalleles (*TtRec8-A1* and *TaRec8-D1*) could independently encode enough Rec8 cohesin essential for normal meiosis in polyploid wheat. These findings provide novel insight into the cross-genome coordination of the *Rec8* homoeologous genes in polyploid wheat, and will facilitate further studies of the homoeologous genes in wheat and other polyploids.

The predicted proteins of *TtRec8-A1, TtRec8-B1*, and *TaRec8-D1* differ from each other at 23 amino acid positions. These amino acid differences separate *TtRec8-B1* from *TtRec8-A1*/*TaRec8-D1*, and *TtRec8-A1* from *TtRec8-B1*/*TaRec8-D1*, suggesting the uniqueness of *TtRec8-B1* and *TtRec8-A1* in polyploid wheat. The protein sequence-based phylogenetic analysis indicates that TtRec8-A1 and TaRec8-D1 are more closely related than their relationship with TtRec8-B1. The protein sequence differences among the *Rec8* homoeoalleles might affect their activity as Rec8 cohesin, and consequently lead to a dominant expression of *TtRec8-A1*/*TaRec8-D1* over *TtRec8-B1* in polyploid wheat.

The differential expression pattern and structural variation of the *Rec8* homoeoalleles might be an evolutionary consequence of the allopolyploid genome in wheat (Zhang *et al.*, 2011). *T. urartu* (2n=2x=14, AA) contributed the A genome to tetraploid wheat (2n=4x=28, AABB) by hybridizing to the B genome ancestor that remains unknown (Dvorak *et al.*, 1993). *Ae. tauschii* (2n=2x=14, DD) contributed the D genome to hexaploid wheat by hybridizing to tetraploid wheat (Kihara, 1944; McFadden and Sears, 1946). Our results and the previous reports consistently indicate that the B genome of tetraploid and hexaploid wheat have significantly higher genetic diversity than the A and D genomes (Chao *et al.*, 1989; Felsenburg *et al.*, 1991; Siedler *et al.*, 1994; Petersen *et al.*, 2006). It appears that the wheat B genome had undergone a more divergent evolutionary process than the A and D genomes (Zhang *et al.*, 2017). In other words, the integrity of the A and D genomes have been well maintained over the evolutionary process, but not quite well for the B genome. As a result, the homoeoalleles in the A and D genomes play a more significant role than those in the B genome, especially for the genes critical in plant development and reproduction, such as *Rec8*. Therefore, the *Rec8* homoeoalleles on chromosomes 1A and 1D may be evolutionarily dominant over the one on chromosome 1B for Rec8 cohesin production in polyploid wheat. Moreover, the differentiation of the *Rec8* homoeoallele on chromosomes 1B from those on chromosomes 1A and 1D implies a distinct origin and evolutionary route of the wheat B subgenome, which ancestor remains unknown, from the A and D subgenomes. These new findings will be beneficial to future genome and evolutionary studies in wheat and its relatives, especially for the B subgenome.

## ACCESSION NUMBERS

Sequence data from this article can be found in the GenBank data libraries under accession numbers MG372313 - MG372315. Sequence data of the *Rec8* orthologues in other species can be found in the GenBank database under the accession numbers listed in the Tables S1 & S3.

## ACKNOWLEDGEMENTS

We thank Wayne Sargent in the Biological Science Lab, USDA-ARS, Fargo, ND for his technical assistance to this research. Also, we would like to thank Dr. Chung-Ju Rachel Wang (Institute of Plant and Microbial Biology, Academia Sinica, Taiwan) and Dr. Wojciech Pawlowski (Cornell University) for their kind help in the immunolocalization procedure. Also, we would like to thank Dr. Rebekah Oliver for critical review of this manuscript. This research was supported by the National Science Foundation under Grant No. 0457356.

## SUPPLEMENTARY DATA

**Figure S1.** Comparative analysis of the cohesin-like proteins.

**Figure S2.** Identification of the BAC clones containing the homoeoalleles *TtRec8-A1* and

*TtRec8-B1* in LDN.

**Figure S3.** CDS alignment of the homoeoalleles *TtRec8-A1, TtRec8-B1*, and *TaRec8-D1*.

**Figure S4.** Predicted protein sequences of the wheat *Rec8* homoeoalleles.

**Figure S5.** Verification of polypeptide pGEX-R26 by protein identification assay.

**Table S1.** Comparative analysis of the predicted TtRec8-A1 protein with the cohesion proteins from other eukaryotic species.

**Table S2.** DNA primers and their sequences used in this study.

**Table S3.** Genbank accession numbers of the Rec8 orthologues involved in the phylogenetic analysis.

## REFERENCES

Bai X, Peirson BN, Dong F, Xue C, Makaroff CA. 1999. Isolation and characterization of SYN1, a RAD21-like gene essential for meiosis in Arabidopsis. Plant Cell 11, 417–430.

Bass HW, Marshall WF, Sedat JW, Agard DA, Cande WZ. 1997. Telomeres cluster de novo before the initiation of synapsis: a three-dimensional spatial analysis of telomere positions before and during meiotic prophase. Journal of Cell Biology 137, 5–18.

Bhatt AM, Lister C, Page T, Fransz P, Findlay K, Jones GH, Dickinson HG, Dean C. 1999. The DIF1 gene of Arabidopsis is required for meiotic chromosome segregation and belongs to the REC8/RAD21 cohesin gene family. Plant Journal 19, 463–472.

Brenchley R, Spannagl M, Pfeifer M, et al. 2012. Analysis of the bread wheat genome using whole-genome shotgun sequencing. Nature 491, 705–710.

Cai X. 1994. Chromosome translocations in the common wheat variety ‘Amigo’. Hereditas 121, 199–202.

Cai X, Dong F, Edelmann RE, Makaroff CA. 2003. The Arabidopsis SYN1 cohesin protein is required for sister chromatid arm cohesion and homologous chromosome pairing. Journal of Cell Science 116, 2999–3007.

Cai X, Xu SS, Zhu X. 2010. Mechanism of ploidy-dependent unreductional meiotic cell division in polyploid wheat. Chromosoma 119, 275–285.

Chao S, Sharp PJ, Worland AJ, Koebner RMD, Gale MD. 1989. RFLP-based genetic maps of homoeologous group 7 chromosomes. Theoretical and Applied Genetics 78, 495–504.

Chao WS, Serpe MD, Jia J, Shelver WL, Anderson JV, Umeda M. 2007. Potential roles for autophosphorylation, kinase activity, and abundance of a CDK-activating kinase (Ea;CDKF;1) during growth in leafy spurge. Plant Molecular Biology 63, 365–379.

Chao WS. 2008. Real-time PCR as a tool to study weed biology. Weed Science 56, 290–296.

Chao WS, Serpe MD. 2010. Changes in the expression of carbohydrate metabolism genes during three phases of bud dormancy in leafy spurge. Plant Molecular Biology 73, 227–239.

Chelysheva L, Diallo S, Vezon D, et al. 2005. AtREC8 and AtSCC3 are essential to the monopolar orientation of the kinetochores during meiosis. Journal of Cell Science 118, 4621–4632.

Dvorak J, Diterlizzi P, Zhang HB, Resta P. 1993. The Evolution of polyploid wheats - identification of the A-genome donor species. Genome 36, 21–31.

Faris JD, Haen KM, Gill BS. 2000. Saturation mapping of a gene-rich recombination hot spot region in wheat. Genetics 154, 823–835.

Felsenburg T, Levy AA, Galili G, Feldman M. 1991. Polymorphism of high molecular weight glutenins in wild tetraploid wheat: spatial and temporal variation in a native site. Israel Journal of Botany 40, 451–479.

Gaut BS. 2002. Evolutionary dynamics of grass genomes. New Phytologist 154, 15–28.

Golubovskaya I, Hamant O, Timofejeva L, Wang CJ, Braun D, Meeley R, Cande WZ. (2006). Alleles of afd1 dissect REC8 functions during meiotic prophase I. Journal of Cell Science 119, 3306–3315.

Hu B, Jin J, Guo AY, Zhang H, Luo JC, Gao G. 2015. GSDS 2.0: an upgraded gene feature visualization server. Bioinformatics 31, 1296–1297.

Huang S, Sirikhachornkit A, Su X, Faris J, Gill B, Haselkorn R, Gornicki P. 2002. Genes encoding plastid acetyl-CoA carboxylase and 3-phosphoglycerate kinase of the Triticum/Aegilops complex and the evolutionary history of polyploid wheat. Proceedings of the National Academy of Sciences of the United States of America 99, 8133–8138.

Huo N, Gu YQ, Lazo GR, Vogel JP, Coleman-Derr D, Luo MC, Thilmony R, Garvin DF, Anderson OD. 2006. Construction and characterization of two BAC libraries from Brachypodium distachyon, a new model for grass genomics. Genome 49, 1099–1108.

Ishiguro K, Watanabe Y. 2007. Chromosome cohesion in mitosis and meiosis. Journal of Cell Science 120, 367–369.

Joppa LR, Williams ND. 1988. Langdon durum disomic substitution lines and aneuploid analysis in tetraploid wheat. Genome 30, 222–228.

Kellogg EA. 2001. Evolutionary history of the grasses. Plant Physiology 125, 1198–1205.

Kihara H. 1944. Discovery of the DD-analyser, one of the ancestors of vulgare wheat. Agriculture and Horticulture 19, 889–890.

Kimbara J, Endo TR, Nasuda S. 2004. Characterization of the genes encoding for MAD2 homologues in wheat. Chromosome Research 12, 703–714.

Kleckner N. 1996. Meiosis: how could it work? Proceedings of the National Academy of Sciences of the United States of America 93, 8167–8174.

Konieczny A, Ausubel FM. 1993. A procedure for mapping Arabidopsis mutations using co-dominant ecotype-specific PCR-based markers. Plant Journal 4, 403–410.

Kumar S, Salyan HS, Gupta PK. 2012. Comparative DNA sequence analysis involving wheat, Brachypodium and rice genomes using mapped wheat ESTs. Triticeae Genomics and Genetics 3, 25–37.

Leach LJ, Belfield EJ, Jiang C, Brown C, Mithani A, Harberd NP. 2014. Patterns of homoeologous gene expression shown by RNA sequencing in hexaploid bread wheat. BMC Genomics 15, 276.

Lee JY, Hayashi-Hagihara A, Orr-Weaver TL. 2005. Roles and regulation of the Drosophila centromere cohesion protein MEI-S332 family. Philosophical Transactions of the Royal Society of London. Series B, Biological Sciences 360, 543–552.

Long YM, Chao WS, Ma GJ, Xu SS, Qi LL. 2017. An innovative SNP Genotyping method adapting to multiple platforms and throughputs. Theoretical and Applied Genetics 130, 597–607.

Ma GJ, Zhang TZ, Guo WZ. 2006. Cloning and characterization of cotton GhBG gene encoding ß-glucosidase. DNA Sequence 17, 355–362.

McFadden ES, Sears ER. 1946. The origin of Triticum speltoides and its free-threshing hexaploid relatives. Journal of Heredity 37, 107–116.

Murai J, Taira T, Ohta D. 1999. Isolation and characterization of the three Waxy genes encoding the granule-bound starch synthase in hexaploid wheat. Gene 234, 71–79.

Murashige T, Skoog F. 1962. A revised medium for rapid growth and bioassays with tobacco culture. Physiologia Plantarum 15, 473–497.

Pearce S, Vazquez-Gross H, Herin SY, Hane D, Wang Y, Gu YQ, Dubcovsky J. 2015. WheatExp: an RNA-seq expression database for polyploid wheat. BMC Plant Biology 15, 299.

Petersen G, Seberg O, Yde M, Berthelsen K. 2006. Phylogenetic relationships of Triticum and Aegilops and evidence for the origin of the A, B, and D genomes of common wheat (Triticum aestivum). Molecular Phylogenetics and Evolution 39, 70–82.

Qi LL, Ma GJ, Long YM, Hulke BS, Gong L, Markell SG. 2015. Relocation of a rust resistance gene R2 and its marker-assisted gene pyramiding in confection sunflower (Helianthus annuus L.). Theoretical and Applied Genetics 128, 477–488.

Roeder GS. 1997. Meiotic chromosomes: it takes two to tango. Genes & Development 11, 2600–2621.

Salse J, Bolot S, Throude M, Jouffe V, Piegu B, Quraishi UM, Calcagno T, Cooke R, Delseny M, Feuillet C. 2008. Identification and characterization of shared duplications between rice and wheat provide new insight into grass genome evolution. Plant Cell 20, 11–24.

Shao T, Tang D, Wang K, Wang M, Che L, Qin B, Yu H, Li M, Gu M, Cheng Z. 2011. OsREC8 is essential for chromatid cohesion and metaphase I monopolar orientation in rice meiosis. Plant Physiology 156, 1386–1396.

Siedler H, Messmer MM, Schachermayr GM, Winzeler H, Winzeler M, Keller B. 1994. Genetic diversity in European wheat and spelt breeding material based on RFLP data. Theoretical and Applied Genetics 88, 994–1003.

Tamura K, Stecher G, Peterson D, Filipski A, Kumar S. 2013. MEGA6: Molecular Evolutionary Genetics Analysis version 6.0. Molecular Biology and Evolution 30, 2725–2729.

Tóth A, Rabitsch KP, Gálová M, Schleiffer A, Buonomo SBC, Nasmyth K. 2000. Functional genomics identifies monopolin: a kinetochore protein required for segregation of homologs during meiosis I. Cell 103, 1155–1168.

Watanabe Y. 2004. Modifying sister chromatid cohesion for meiosis. Journal of Cell Science 117, 4017–4023.

Watanabe Y. 2005. Shugoshin: guardian spirit at the centromere. Current Opinion in Cell Biology 17, 590–595.

Watanabe Y. 2012. Geometry and force behind kinetochore orientation: lessons from meiosis. Nature Reviews Molecular Cell Bioogy 13, 370–382.

Watanabe Y, Nurse P. 1999. Cohesin Rec8 is required for reductional chromosome segregation at meiosis. Nature 400, 461–464.

Yokobayashi S, Watanabe Y. 2005. The kinetochore protein Moa1 enables cohesion-mediated monopolar attachment at meiosis I. Cell 123, 803–817.

Yu HG, Dawe RK. 2000. Functional redundancy in the maize meiotic kinetochore. Journal of Cell Biology 151, 131–141.

Yuan L, Yang X, Ellis JL, Fisher NM, Makaroff CA. 2012. The Arabidopsis SYN3 cohesin protein is important for early meiotic events. Plant Journal 71, 147–160.

Zhang L, Tao J, Wang S, Chong K, Wang T. 2006. The rice OsRad21-4, an orthologue of yeast Rec8 protein, is required for efficient meiosis. Plant Molecular Biology 60, 533–554.

Zhang W, Zhang M, Zhu X, Cao Y, Sun Q, Ma G, Chao S, Yan C, Xu S, Cai X. 2018. Molecular cytogenetic and genomic analyses reveal new insights into the origin of the wheat B genome. Theoretical and Applied Genetics 131, 365–375.

Zhang Z, Belcram H, Gornicki P, et al. 2011. Duplication and partitioning in evolution and function of homoeologous *Q* loci governing domestication characters in polyploid wheat. Proceedings of the National Academy of Sciences of the United States of America 108, 18737–18742.

